# Elevated serum adenosine deaminase levels in neuroleptic-naïve patients with recent-onset schizophrenia

**DOI:** 10.1101/070748

**Authors:** Arun Sasidharan, Sunil Kumar, John P John, Mariamma Philip, Sarada Subramanian, Sanjeev Jain, Bindu M Kutty

## Abstract

Schizophrenia is characterized by pathophysiological alterations of multiple neurotransmitter systems such as dopaminergic, glutamatergic, GABA-ergic and serotonergic pathways. Adenosine, a homeostatic neuromodulator that mediates signaling through multiple neurotransmitter pathways, is an emerging candidate neurobiological substrate of schizophrenia. The present study examined peripheral blood levels of adenosine deaminase, an adenosine metabolizing enzyme, in 16 neuroleptic-naive patients with recent-onset schizophrenia (mean age=25.59 years (range: 16-35)) and 18 age-matched healthy comparison subjects (mean age=25.17 years (range: 18-28)). Serum adenosine deaminase levels were assayed at two time points; before (7 p.m.) and after (7 a.m.) sleep. The adenosine deaminase levels were compared between groups and were correlated to positive and negative symptom severity measures. Adenosine deaminase levels were found to be higher at both evening (*p*=0.013) and morning (*p*<0.001) time points in our sample of patients with recent-onset schizophrenia who were never exposed to neuroleptic medications. Correlational analysis revealed evidence for a possible link between evening rise in adenosine deaminase and severity of auditory hallucinations (*p*=0.003) as well as morning rise in adenosine deaminase and severity of avolition-apathy in patients with schizophrenia (*p*=0.013). The results of the study provide strong support to the adenosine hypothesis of schizophrenia and highlight the potential utility of serum adenosine deaminase as a peripheral biomarker of schizophrenia.

## 1. Introduction

The pathophysiology of schizophrenia is understood to involve dysfunction of the dopaminergic (Carlsson, 1988), serotonergic (Meltzer and Massey, 2011), glutamatergic (Coyle, 2006) and GABA-ergic (Gonzalez-Burgos and Lewis, 2008) neurotransmitter systems. The diverse psychopathology of schizophrenia cannot be explained by dysfunction of neurotransmitter systems when considered in isolation (Keshavan et al., 2011). Adenosine, a homeostatic neuromodulator, is an emerging candidate neurobiological substrate with effects on multiple neurotransmitter pathways (Boison, 2008; Cunha and Cunha, 2001). In addition, adenosine plays an important role in early brain development and regulation of brain immune responses (Cunha and Cunha, 2001), and thereby could also contribute to the neurodevelopmental deviations implicated in schizophrenia (Lara et al., 2006).

Adenosine deaminase (ADA) is a purine-inactivating endoenzyme that irreversibly deaminates adenosine to inosine (Yegutkin, 2008), leading to its final degradation to uric acid. Like adenosine, ADA too is ubiquitously found in the human body (Franco et al., 1997) and hence implicated in diverse physiological functions. Thus, serum ADA level has been suggested as an important peripheral biomarker of adenosine signaling in neuropsychiatric disorders (Elgün et al., 1999; Herken et al., 2007; Stubbs et al., 1982) especially in schizophrenia, given that a hypoadenosinergic state has been linked to its pathogenesis (Lara et al., 2006).

In support of the above, Dutra et al. (Dutra et al., 2010) demonstrated lower frequency of occurrence of an ADA variant with decreased enzymatic activity (G/A genotype) among patients with schizophrenia, suggesting increased levels of ADA and reduced levels of ambient adenosine. Another study reported significantly higher serum ADA in patients with schizophrenia on antipsychotic monotherapy (more so with atypical antipsychotic), when compared to healthy control subjects (Brunstein et al., 2007), but found no correlation with clinical psychopathology. However, Ghaleiha et al. (Ghaleiha et al., 2011), reported that in patients with chronic schizophrenia, antipsychotic therapy (particularly with clozapine) was associated with an increase in serum ADA and symptomatic improvement. Therefore, it is unclear whether the increased serum ADA reported in the above studies was the consequence of treatment with antipsychotics or a marker of the disorder per se.

Therefore, we examined whether serum ADA levels were significantly different in patients with schizophrenia who have never been exposed to antipsychotic medications in comparison to matched healthy comparison subjects. In accordance with the proposed adenosine theory of schizophrenia pathophysiology, we hypothesized that the serum ADA will be significantly higher in patients with schizophrenia. We also aimed at exploring the hitherto unreported relationship between serum ADA and symptom severity scores.

## 2. Materials and Methods

The study was carried out at the National Institute of Mental Health and Neurosciences (NIMHANS), Bangalore, India, with due approval from the Institute Ethics Committee thus conforming to the ethical standards laid down in the 1964 Declaration of Helsinki. Written informed consent was obtained from all the participants prior to enrolment into the study.

### 2.1. Participants

20 patients with schizophrenia (SZ) or schizophreniform disorder and 20 age-matched healthy comparison subjects (HS) participated in the study. Of the above subjects, the blood samples of four SZ and two HS were of inadequate quantity and/or poor quality, and therefore had to be omitted from the study. The remaining participants (SZ=16; HS=18) were of similar age group (SZ: mean=25.59 years (range: 16-35); HS: mean=25.17 years (range: 18-28)). Patients with schizophrenia/schizophreniform disorder had a mean illness duration of 21 months (range: 1-96). Of these, 11 SZ and 17 HS had ADA samples collected at two time points, i.e., at 7p.m. on day1 and 7a.m. on day2. Five SZ had only their morning samples collected, since they did not abstain from consuming food for 3 hours prior to the evening sample collection. The sample collection could not be rescheduled to the next day for ethical reasons, since the patients were neuroleptic naïve and had to be started on medications at the earliest. In one of the healthy subjects also, the morning sample alone could be collected due to certain logistic constraints.

The diagnosis of schizophrenia/schizophreniform disorder was arrived at using criteria from the Diagnostic and Statistical Manual for Mental Disorders – Fourth Edition (DSM-IV) (American Psychiatric Association, 2000) based on the consensus of a research psychiatrist who conducted a semi-structured interview and a trained research assistant who used the Mini International Neuropsychiatric Interview (MINI) for DSM-IV (Sheehan et al., 1998). Positive and negative symptoms were rated using the Scale for Assessment of Positive symptoms (SAPS) (Andreasen, 1983) and Scale for Assessment of Negative symptoms (SANS) (Andreasen, 1982) respectively by one rater (S.K.) for all subjects after undergoing adequate training and establishment of inter-rater reliability. None of the patients had history of exposure to neuroleptic medications.

The healthy comparison subjects were recruited from the community through word-of-mouth. They were screened to exclude past history of Axis I psychiatric disorders, personal history of psychoactive medication use and family history of schizophrenia spectrum disorders in first-degree relatives using a study-specific proforma.

All participants were screened to exclude history of significant head injury, neurological disorders, medical conditions including acute infections, autoimmune disorders, endocrine disorders, specific sleep disorders, and substance abuse including caffeine (daily intake of caffeine-containing beverages and food substances exceeding 300mg/day). All were free of any drugs known to affect immune or endocrine function. All participants underwent clinical screening to rule out any unstable medical conditions. The participants were instructed to avoid food and caffeine for at least 3 hours before the evening blood sample, following which they had their dinner; overnight fasting was ensured before drawing the morning blood sample. The above precautionary measures were adopted to avoid any confounding effects, including that of food and caffeine on the measured ADA levels.

### 2.2. Measurement of ADA

Blood samples were collected from participants before and after their sleep as there was no available evidence from the literature regarding the most appropriate time to collect serum for ADA assay. The first sample was collected around 7p.m. (before dinner and before sleep) similar to a previous schizophrenia study (Brunstein et al., 2007) and the second sample around 7a.m. the following morning (after sleep and before breakfast) similar to earlier studies in major depression (Elgün et al., 1999; Herken et al., 2007). The participants slept in the sleep cabin of the Sleep Research Laboratory at the Department of Neurophysiology. 5ml of venous blood samples were collected in vacuum tubes (BD Vacutainer®) without anticoagulants on both occasions. The 7p.m. samples were allowed to clot overnight at 4°Celsius whereas the 7a.m. samples were allowed to clot at room temperature for two hours. Later both the samples were centrifuged for 10 minutes at 4000rpm to extract the serum. The serum samples were then coded and stored in microtubes (Eppendrof Inc.) at -80°Celsius for the assay.

The ADA assay from serum was performed using a previously validated (Al-Rubaye and Morad, 2012, 2013) colorimetric sandwich-enzyme immunoassay kit E91390Hu96 (USCN Life Science Inc., Wuhan), following the instructions provided in the manual (“SEB390Hu-96 ELISA Kit for Adenosine Deaminase (ADA) - Instruction manual,” 2012). All samples were analyzed in duplicate and the intra-assay variability was less than 10%. Assays were conducted at the Department of Neurochemistry under the supervision of S.S. The trained neurochemist who carried out the assays was blinded to the study group status of the blood samples.

### 2.3. Genotyping of ADA polymorphism

The ADA 22G>A polymorphism (rs73598374) was genotyped using allele-specific polymerase-chain reaction under the supervision of M.P and S.J. Genotyping was done for 32 (HS=16; SZ=16) out of the 35 subjects, all of whom were found to have the G/G genotype. Therefore, any further examination of the relationship between the ADA rs73598374 genotypes and serum ADA levels and the differences between patients with schizophrenia and healthy comparison subjects was not possible in our limited sample.

### 2.4. Statistical analysis

Statistical analysis was performed using GraphPad Prism 5.0 (GraphPad Software, Inc.) and Statistical Toolbox of MATLAB 2012b (Mathworks, USA). D'Agostino and Pearson omnibus test and visual inspection of histogram plot was used to determine the normality of data distribution. As ADA levels of both groups did not follow normal distribution, statistical analyses were done on their log transformed values which followed a normal distribution. All other values, except age, followed normal distribution. Mann–Whitney U-test was used to test the group difference in age. Between-group comparisons for ADA levels of the 7p.m. (ADA_even_) and of the 7a.m. (ADA_morn_) samples as well as their difference (ADA_diff(even-morn)_), were done using the Student's t-test, after conducting a permutation-based two-way ANOVA that found no group vs time interactions (Table 1). Comparisons between ADA_even_ and ADA_morn_ levels within each group were carried out using paired t-test. The significance level for these tests was set at p<0.05. Pearson's test was used to test correlations of ADA levels (ADA_even_, ADA_morn_ and ADA_diff(even-morn)_) with duration of illness, positive symptom scores and negative symptom scores within the schizophrenia group. Spearman rank order correlation test was used for the within-group correlations between ADA levels and age. All the SAPS and SANS sub-scores (including their total scores) were included in the correlation analysis, and thus 11 correlations were made for each ADA level (ADA_even_, ADA_morn_ and ADA_diff(even-morn)_). To exclude accidental significance associated with multiple correlations, a threshold value of 0.735 was set for the absolute correlation co-efficients (α=0.05; n=11) using G*power 3.1 software (Faul et al., 2009, 2007) in order to obtain a statistical power greater than 80% (Sabri et al., 1997). All tests were assessed using two-tailed p values.

**Table 1:** Two-way ANOVA of ADA levels between groups and sample collection times

|  | CNT (n=17<br>pairs) | SCZ (n=11<br>pairs) |  |  |
| --- | --- | --- | --- | --- |
| Evening<br>sample | 0.862 (0.280) | 1.129 (0.227) | F=-<br>2.782;<br>p=0.009 | F=-28.671;<br>p<0.001 |
| Morning<br>sample | 0.731 (0.240) | 1.111 (0.243) | F=-<br>3.419;<br>p=0.002 |  |
|  | F=0.787;p=0.432 | F=0.181;p=0.853 | F=0.226;p=0.699 |  |
|  | F=0.650;p=0.528 |  |  |  |

**NOTE:** Permutation-based two-way ANOVA was used with 10,000 re-samples on log-transformed serum ADA levels. Mean (SD) values are given in the central while cells. Outer dark grey cells show the F- and p-values. Note that significant main effects exist only for group factor, and no interaction effect is seen. CNT – healthy controls; SCZ – patients with schizophrenia.

## 3. Results

In the two-way ANOVA, significant main effects were noted only for the group factor, while there were no time effects or group × time interactions (Table 1). On further exploring the group difference, patients with schizophrenia showed significantly higher ADA_even_ (Fig.1A; t=2.66, p=0.013) and ADA_morn_ (Fig.1B; t=3.79, p<0.001) levels, when compared to healthy comparison subjects. ADA_even_ and ADA_morn_ showed significant correlation in both HS (n=17pairs, r=0.706, p=0.002) and SZ (n=11pairs, r=0.760, p=0.007) groups. ADA_even_ appeared to be higher than ADA_morn_ in both SZ (meanADA_even_=15.38ng/mL, SD=9.13, n=11; meanADA_morn_=14.08ng/mL, SD=5, n=16), and HS (meanADA_even_=8.85 ng/mL, SD=6.10, n=17; meanADA_morn_=7.16ng/mL; SD=4.18, n=18). But the paired t-test did not find significant difference between ADA_even_ and ADA_morn_ in both HS (t=1.43, p=0.1712) and SZ (t=0.368, p=0.7203). We performed correlational analyses between ADA levels and SANS and SAPS sub-scores to explore the relationship, if any, between symptom severity and ADA levels (Table 2). A significant positive correlation was noted between ADA_diff(even-morn)_ and hallucination sub-score of SAPS (Fig.2B; n=11pairs, r=0.810, p=0.003) at the a-priori-decided significance threshold to account for multiple comparisons (see Statistical analysis subsection). However, there was a high negative correlation nearing significance between ADA_diff(even-morn)_ and avolition-apathy sub-score of SANS (Fig.2C; n=11pairs, r=-0.717, p=0.013). A positive trend was observed between ADA_even_ and hallucination sub-score of SAPS (n=11pairs, r=0.601, p=0.050) while ADA_morn_ showed a trend for a positive correlation with avolition-apathy sub-score of SANS (n=16pairs, r=0.500, p=0.049).

**Fig. 1:**
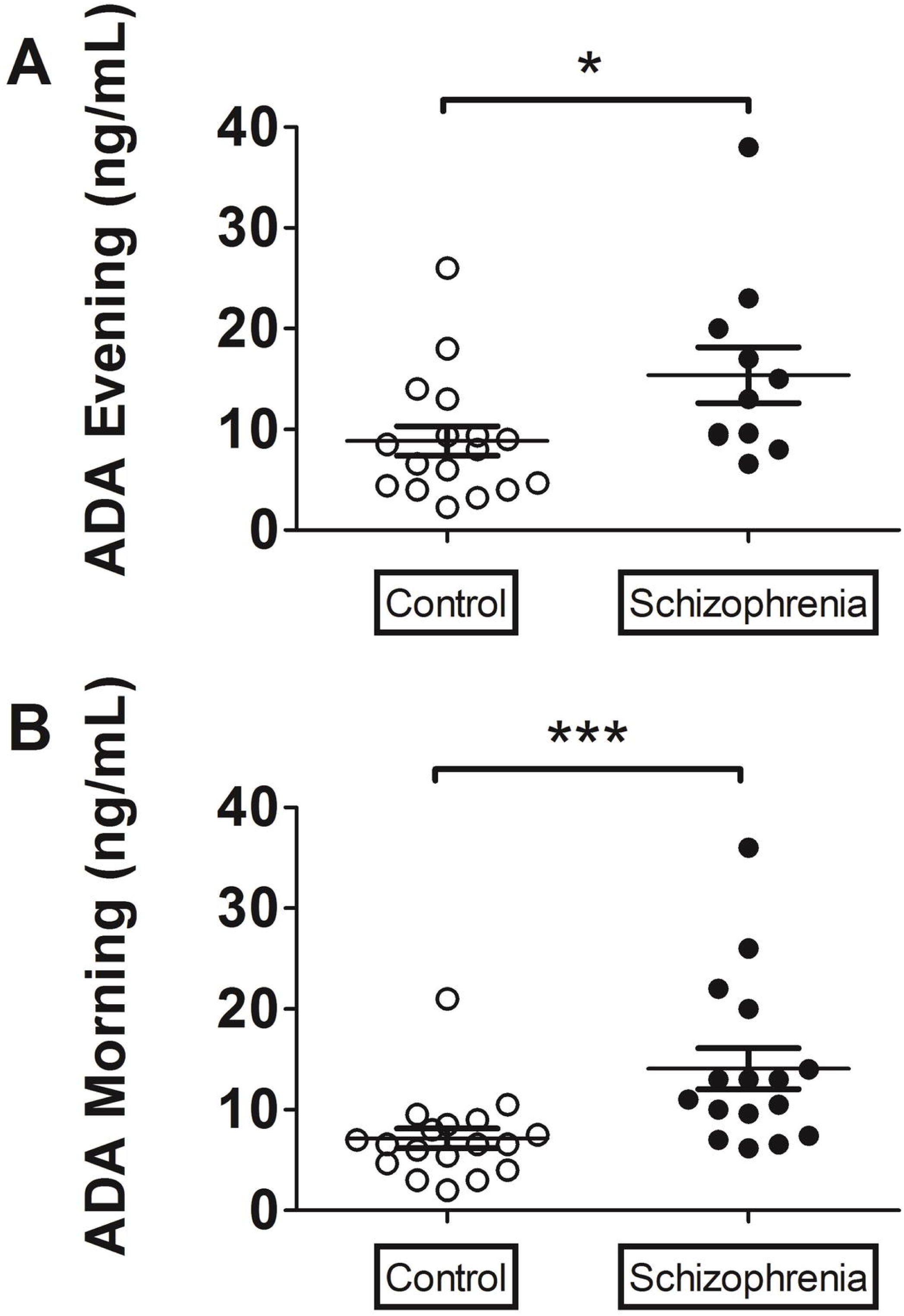
Serum levels of Adenosine deaminase (ADA). Scatter plots showing serum ADA levels in: **A)** evening sample [control=17; schizophrenia=11], and **B)** morning sample [control=18; schizophrenia=16]. Unpaired t test performed between log transformed ADA values. **\*p < 0.05; ***p < 0.001**.

**Fig. 2:**
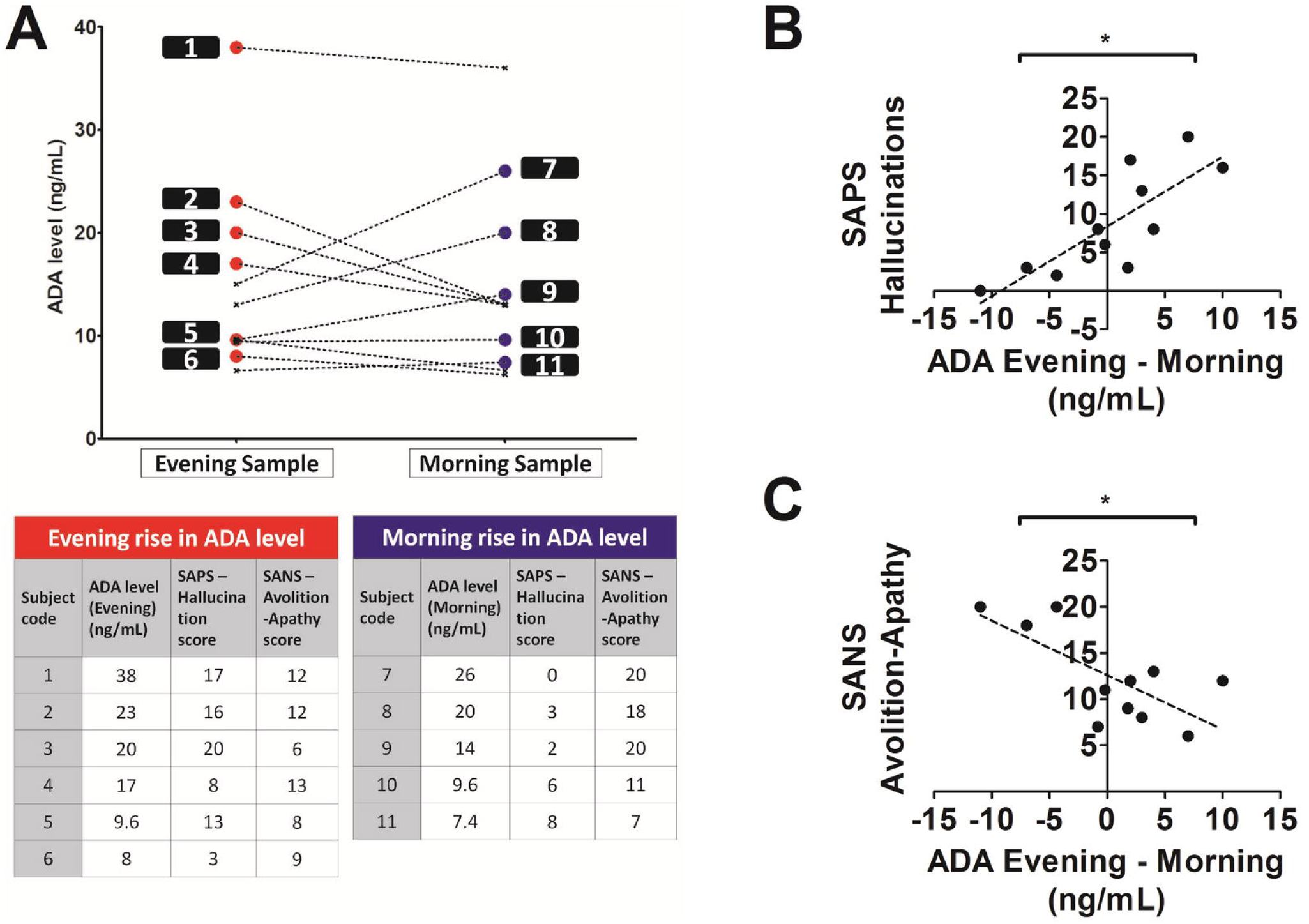
Relation between Adenosine deaminase (ADA) levels and symptom severity. **A)** Line graph and table showing the relation between relatively higher ADA levels in the evening samples (evening rise in ADA) or in the morning samples (morning rise in ADA), with symptom severity of patients with schizophrenia [n=11]. Note that patients with higher evening rise in ADA (shown in red) have higher hallucination scores, while those with higher morning rise in ADA (shown in blue) have higher avolition-apathy scores. Line graph showing the correlation of difference between evening and morning ADA levels versus: **B)** Scale for Assessment of Positive symptoms (SAPS) hallucination sub-score [n=11pairs, r=0.8097, p=0.0025], and **C)** Scale for Assessment of Negative symptoms (SANS) avolition-apathy sub-score [n=11pairs, r=-0.7165, p=0.0131]. Pearson's correlation test (two-tailed). **\*r > 0.735 (see Statistical Analysis section)**.

**Table 2:** Correlational analysis between ADA levels and symptom severity.

|  | Scale for Assessment of Positive symptoms<br>(SAPS) |  |  |  |  | Scale for Assessment of Negative symptoms<br>(SANS) |  |  |  |  |  |
| --- | --- | --- | --- | --- | --- | --- | --- | --- | --- | --- | --- |
|  | Hallucinations | Delusions | Bizarre behaviour | Positive formal thought disorder | Total score | Affective flattening | Alogia | Avolition-apathy | Anhedonia | Attention | Total score |
| **ADA**<br>**(Evening – Morning)** | **0.81**<br>**(0.003)** | 0.18<br>(0.591) | -0.64<br>(0.035) | -0.38<br>(0.250) | 0.22<br>(0.506) | -0.17<br>(0.623) | -0.39<br>(0.240) | **-0.72**<br>**(0.013)** | -0.49<br>(0.130) | -0.08<br>(0.819) | -0.41<br>(0.206) |
| **ADA**<br>**(Evening)** | 0.60<br>(0.050) | 0.19<br>(0.580) | -0.38<br>(0.253) | -0.25<br>(0.457) | 0.21<br>(0.530) | -0.09<br>(0.796) | -0.09<br>(0.787) | 0.10<br>(0.767) | -0.20<br>(0.547) | 0.10<br>(0.777) | -0.07<br>(0.839) |
| **ADA**<br>**(Morning)** | 0.05<br>(0.878) | 0.10<br>(0.772) | 0.02<br>(0.959) | 0.04<br>(0.917) | 0.08<br>(0.807) | 0.07<br>(0.837) | 0.20<br>(0.546) | **0.60**<br>**(0.049)** | 0.17<br>(0.623) | 0.14<br>(0.673) | 0.25<br>(0.464) |
| **NOTE:** Pearson's Correlation test (two-tailed) was performed between log-transformed ADA levels and scores of SAPS and SANS. All data shown as: Correlation coefficient (p-value). Significant correlation is shown in bold and highlighted in dark grey. Trend correlations are shown in italics and highlighted in light grey. |  |  |  |  |  |  |  |  |  |  |  |

ADA levels were not found to be significantly correlated with age in HS (ADA_even_: n=17pairs, r=-0.092, p=0.727; ADA_morn_: n=18pairs, r=-0.363, p=0.139) and in SZ (ADA_even_: n=11pairs, r=0.097, p=0.777; ADA_morn_: n=16pairs, r=-0.190, p=0.481). ADA levels were also not significantly correlated with duration of illness in the SZ (ADA_even_: n=11pairs, r=-0.044, p=0.897; ADA_morn_: n=16pairs, r=-0.047, p=0.862).

## 4. Discussion

The present study found significantly higher serum ADA levels at two different time points (7p.m. on day1 and 7a.m. on day2) in patients with recent-onset schizophrenia who had never been exposed to neuroleptic medications, in comparison to matched healthy comparison subjects. The study also provides evidence for a link between ADA levels and psychopathology in patients with schizophrenia.

As described in the Introduction, similar observations have been reported in medicated patients with chronic schizophrenia (Brunstein et al., 2007; Ghaleiha et al., 2011). The results of our study on neuroleptic-naïve patients with recent-onset schizophrenia provide the initial evidence towards considering increased serum ADA as a potential peripheral marker of a hypoadenosinergic state in schizophrenia (Lara et al., 2006), and not just a secondary effect of illness chronicity or neuroleptic treatment (Brunstein et al., 2007; Ghaleiha et al., 2011).

The higher ADA levels in patients would imply faster and greater degradation of adenosine, leading to a hypoadenosinergic state. Adenosine inhibits the release of several neurotransmitters, such as glutamate, dopamine, serotonin and acetylcholine, and decreases neuronal activity by post-synaptic hyperpolarization, thereby serving an important neuromodulatory function (Lara et al., 2006). Animal models with altered adenosine signaling have provided important insights into the adenosine dependent changes in neurotransmitter signaling associated with the genesis of schizophrenia (Ferré, 1997; Lara, 2002; Sills et al., 1999). Thus the increased ADA levels observed in our study could be the marker of an adenosine deficit state, which results in impaired neuromodulation in multiple brain networks leading on to the various symptoms of schizophrenia. Further support to the adenosine hypothesis of schizophrenia comes from a genetic study (Dutra et al., 2010) that has reported a lower proportion of ADA genotype with reduced activity (G/A) among patients with schizophrenia.

Though, as a group, patients with schizophrenia had significantly higher ADA at both time points, there was indeed wide variability of ADA values within and between the two time points (Fig. 1B). This might reflect the within-group differences in severity of the various psychopathological dimensions. Interestingly we found a significant positive correlation between ADA_diff(even-morn)_ and auditory hallucinations and a trend towards an inverse correlation between ADA_diff(even-morn)_ and avolition-apathy. The positive correlation between higher ADA_even_ (relative to ADA_morn_) and auditory hallucinations (Fig. 2A and 2B) might reflect the inhibitory deficit in the fronto-temporo-parietal networks associated with hyperglutamatergia (Schobel et al., 2013; Théberge et al., 2002) and hyperdopaminergia (Heinz and Schlagenhauf, 2010) secondary to a hypoadensoinergic state (Lara et al., 2006) during the day. Similarly, the positive correlation between higher ADA_morn_ (relative to ADA_even_) and avolition-apathy (Fig. 2A and 2C) might reflect the hypoadenosinergic state during night causing disrupted sleep, leading on to lethargy and avolition during day time. These findings would therefore provide additional support to the possibility of a hypoadenosinergic state (Boison, 2011; Lara et al., 2006) mediating the predominant symptom dimensions of schizophrenia.

As mentioned in the Introduction, adenosine deaminase (ADA) is primarily responsible for catalyzing the irreversible deamination of adenosine to inosine. At least two isoenzymes of ADA viz., ADA1 and ADA2 are identified in humans (Ratech et al., 1981). While these isoenzymes are present in several tissues, they differ in their kinetic properties and tissue distribution. ADA1 is mostly intracellular or on the cell membrane in the ecto-form, attached to dipeptidyl peptidase 4 (Fan et al., 2012). On the other hand, ADA2 is the main isoenzyme in the serum. While ADA1 is expressed in lymphocytes and macrophages, ADA2 is more abundant in the blood, brain and liver (Rosemberg et al., 2007). Previous studies cited earlier (e.g., Brunstein et al., 2007; Ghaleiha et al., 2011) have also assayed serum levels of ADA as a surrogate marker for ADA levels in the brain.

It is pertinent to mention at this juncture that adenosine and other nucleosides are transported across the blood-brain barrier via a saturable, carrier mediated mechanism (Kalaria and Harik, 1988). The increased ADA activity in the periphery, as seen in patients with drug naïve schizophrenia, may result in reduction of the circulatory levels of adenosine. This may decrease the transport across the blood-brain barrier (BBB) leading to a hypoadenosinergic state in the CNS. However, this speculation requires confirmation with additional experiments involving the quantitation of adenosine and its metabolites (inosine, hypoxanthine) in serum and CSF by HPLC method and measuring the isoform specific enzyme activity in the serum and CSF.

It may be argued that the higher ADA found in our patient sample may be a compensatory phenomenon secondary to a state-dependent increase in brain adenosine levels during the psychotic state (Cunha and Cunha, 2001; Hirayama et al., 2011; Reddy et al., 1992). However, this argument fails to explain the overall ADA rise noted even among patients with the negative symptom of avolition-apathy. As our patient group comprised of subjects with heterogeneous symptom dimensions (Fig. 2A), it is unlikely that the higher ADA could have exclusively resulted from higher state-dependent adenosine levels. Thus, the increased ADA levels detected in our sample of patients with schizophrenia may be considered as a peripheral marker of a hypoadenosinergic state that characterizes the disorder. Similarly, medication induced elevation of serum ADA reported by Ghaleiha et al. (2011) in patients with schizophrenia could be compensatory to an improvement in adenosinergic tone associated with successful treatment. It has to be stated that these are indeed preliminary hypotheses generated from the results of the study, which need to be tested in future studies with larger sample sizes.

Adenosine has been shown to serve a regulatory function in the immune system of brain (Haskó et al., 2005). Though general immunological disorders were ruled out in all our subjects through careful history and routine clinical hematological and biochemical screening investigations, we did not carry out screening blood investigations to rule out autoimmune conditions. Although remote associations between acute psychotic states, encephalitis and NMDA auto-antibodies have been reported (Deakin et al., 2014), schizophrenia has not been conclusively shown to be an autoimmune disorder (Coutinho et al., 2014). Therefore, it is unlikely that the raised ADA levels observed in our sample of patients with schizophrenia could be secondary to an autoimmune state.

A potential limitation of the study is the fact that only protein levels of ADA in serum were measured, while enzymatic activity of ADA was not assayed. While measuring the enzyme activity offers greater correlation with physiological response, the practical difficulties in handling the samples prompted us to measure the ADA protein levels. In our earlier pilot studies, we observed a gradual decrease in the ADA enzyme activity when the serum samples were stored at 4°C or when subjected to two freeze-thaw cycles. However, as expected, the ADA protein content remained unaltered upon storage/ freeze thawing. The present study involved the recruitment of drug naïve subjects with schizophrenia over a substantial time period. The samples were collected as and when the subjects were recruited and the serum separated and stored. To avoid inter-assay variation, all the samples were analyzed at the same time. In order to avoid any bias in the results due to varying lengths of sample storage, it was considered appropriate to measure the protein content. Moreover, ADA also has significant non-enzymatic actions (protein-protein interactions) with respect to desensitizing and enhancing functionality of adenosine receptors (Ciruela et al., 2010; Gracia et al., 2008). Therefore, a protein level assessment may be a better indicator of the full range of ADA activity in comparison to enzyme activity.

Finally, in our limited sample of neuroleptic-naïve patients with recent-onset schizophrenia and matched healthy comparison subjects, all the genotyped subjects (HS=16; SZ=16) were found to have G/G genotype of the ADA 22G>A polymorphism (rs73598374); none of the subjects had the G/A genotype that is associated with lower ADA activity (Bachmann et al., 2012; Battistuzzi et al., 1981). Larger samples of schizophrenia and healthy subjects may be needed to observe between-group differences, if any, of the G/G and G/A genotypes as reported in one previous study (Dutra et al., 2010). Nevertheless, since all our subjects were homogeneous with respect to the ADA 22G>A functional polymorphic variation, it may be inferred that our observation of higher serum ADA levels in patients with schizophrenia is unlikely to be confounded by genotypic variation between the study samples.

## 5. Conclusion

To the best of our knowledge, this is the first report of elevated serum ADA levels at two time points in neuroleptic-naïve patients with recent-onset schizophrenia. Further, the study also provides preliminary evidence for a link between ADA levels at different time points with positive and negative symptoms of schizophrenia. These findings may be considered strong evidences in support of the adenosine hypothesis of schizophrenia. In the background of previous reports suggesting a possible genetic basis for the elevated ADA activity in schizophrenia and the observation of alteration of ADA levels with successful treatment, the potential utility of serum ADA levels as a biomarker or endophenotype of schizophrenia should be researched in future studies.

## 6. Acknowledgments

We would like to thank Mrs. Ammu Lukose (for recruitment and evaluation of subjects), Dr. Mathew John (for serum extraction and storage), Dr. Meera Purushottam (for genetic analysis), and all the participants who devoted their time and efforts to take part in this study. This work was funded in part by Department of Biotechnology (DBT), Government of India (Grant No. BT/PR/8363/MED/14/1252 & DST/CSI/100/1321 to J.P.J), and Department of Science & Technology (DST), Government of India (Grant No. SR/SO/BB-27/2009 to S.S.).

## 8. Contributors

A.S and S.K. carried out the data acquisition and analysis; A.S., S.K., B.M.K. and J.P.J. conceptualized the study; M.P. helped with statistical analysis; S.S. facilitated the ADA assessments; S.J. facilitated the Genetic assessments; B.M.K. and J.P.J. critically evaluated the study; A.S and J.P.J. wrote the manuscript and all other authors contributed to writing the manuscript.

